# Transcriptome profiling of cell-free RNAs in prostate cancer patients on active surveillance

**DOI:** 10.64898/2026.09.03.744568

**Authors:** John Gordon Foster, Eva Wozniak, Charles A. Mein, James Thompson, Phillip Stricker, Anne-Maree Haynes, Prabhakar Rajan

**Affiliations:** Centre for Cancer Cell and Molecular Biology Barts Cancer Institute, Queen Mary University of London, London, UK; Focyte, 1 Northway, Altrincham, Cheshire, UK; Genome Centre, Queen Mary University of London, London, UK; Department of Urology, St. George Hospital, Kogarah, Australia; St. Vincent’s Prostate Cancer Research Centre, Darlinghurst, Australia; Garvan Institute of Medical Research and Kinghorn Cancer Centre, Darlinghurst, Australia; Division of Surgery and Interventional Sciences, University College London, UK; Department of Urology, Barts Health NHS Trust, London, UK; Department of Urology, University College London Hospitals NHS Foundation Trust, London. UK

## Abstract

In this feasibility study, plasma cfRNA transcriptomes from 64 men with prostate cancer on active surveillance revealed differences between stable disease and progression to radical prostatectomy and suggested a predominantly non-tumour origin of circulating RNA. These findings demonstrate the feasibility of cfRNA transcriptomic analysis in this setting and support its further investigation as a minimally invasive adjunct for personalised risk stratification and monitoring.

## Main Text

Active surveillance (AS) delays or avoids unnecessary treatment-related side effects for early localised non-aggressive prostate cancer (PCa) through monitoring with regular prostate-specific antigen (PSA) testing, digital rectal examination (DRE), magnetic resonance imaging (MRI) and prostate biopsy^[1]^. Whilst baseline MRI has prognostic value and is increasingly used for disease monitoring, molecular biomarkers could be an adjunct to refine risk stratification and disease prognostication^[2]^. Unlike tissue-based genomic classifiers such as Decipher (Veracyte) and Prolaris CCP (Myriad), blood-based molecular testing for example cell-free RNA (cfRNA) offers minimally invasive real-time monitoring^[2,3]^. Moreover, cfRNA may yield genetic information shed from different cell populations within the tumour microenvironment, as well as the host circulation, expanding the opportunities for biomarker discovery. Here, we test the feasibility of whole transcriptome profiling of cfRNAs from the blood of PCa patients under AS, exploring differences between patients with stable disease (SD), and progression to radical prostatectomy (RP).

Sixty-four patients were selected from a cohort of AS patients with routinely collected baseline plasma samples following ethical approval and informed consent, of which 33 had SD and 31 had progressed to RP at the time of last follow up. Plasma cfRNA was extracted and subjected to RNA-sequencing using standard library preparation, sequencing, and computational analysis workflows (Fig. 1A). Most aligned reads mapped to exonic regions following removal of PCR duplicates (Fig. 1B). RP reads were significantly shorter and had lower alignment rates than those from the SD group (Supplementary Fig. 1A-B).

**Fig. 1.**
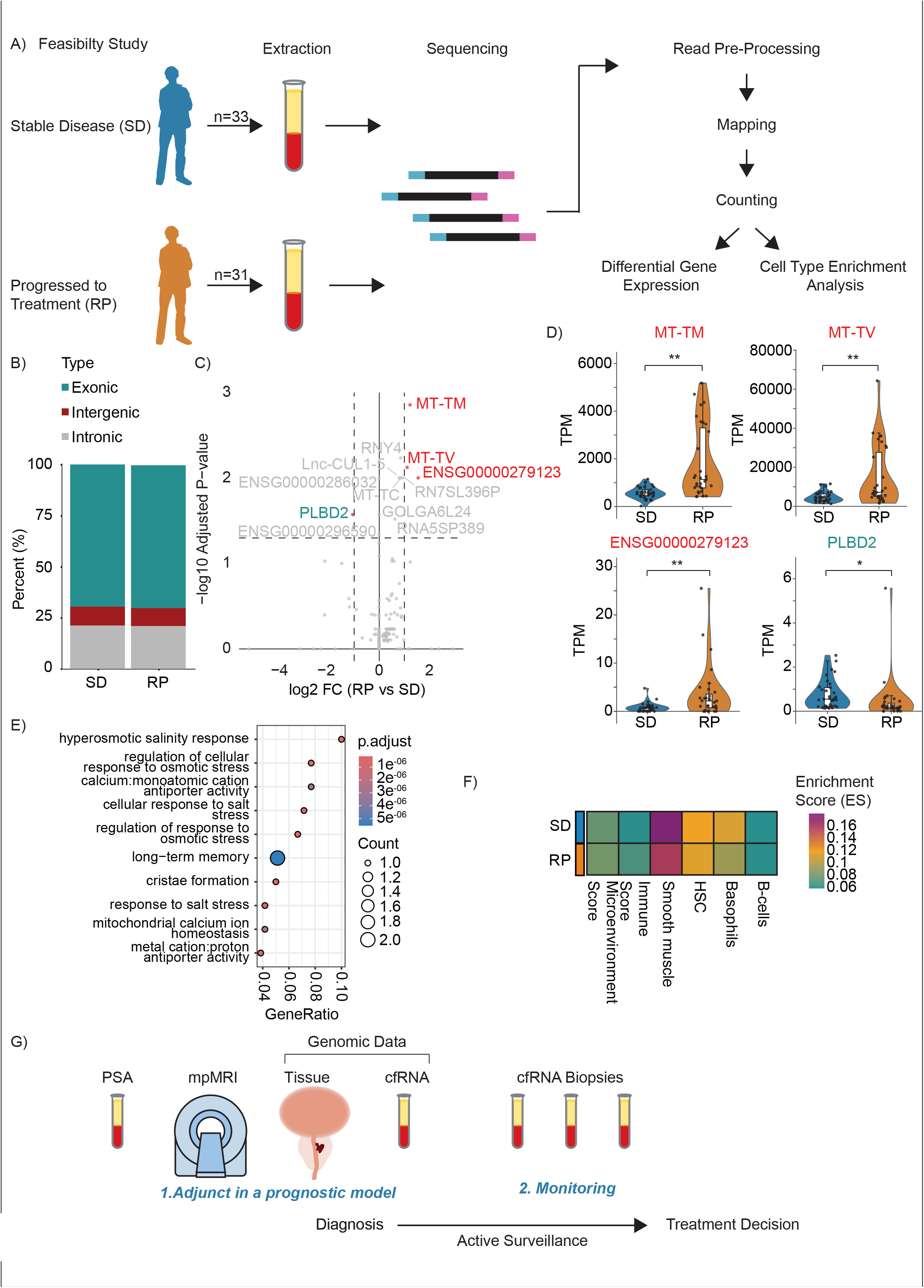
– Feasibility study of cfRNA sequencing in an AS cohort of patients. (A) Schematic representing sample processing and cell free RNA sequencing analysis workflow use in this feasibility study. (B) Stacked bar plot of the genomic region of aligned reads for SD (Stable Disease) and RP (Progressed to Radical Prostatectomy) patients. (C) Volcano plot of differentially expressed genes between RP and SD patients. Coloured genes indicate those with log_2_FC >=1 (red) or =< -1 (teal) and adjusted pvalue <0.05. Named genes in grey are those with adjusted pvalue <0.05 but which do not meet the log2FC threshold. (D) Violin plots representing down-regulated (teal) and up-regulated (red) genes, comparisons are between RP and SD patients, adjusted pvalues from differential gene expression analysis are given * padj <0.05, ** padj <0.01. (E) Gene Ontology analysis using gseGO function of Cluster Profiler in R. (F) Heatmap of xCell cell type enrichment analysis of gene expression matrix (TPM) of RP and SD patients, for categories where enrichment score (ES) > 0.05. (G) cfRNA transcriptomic profiling as a minimally invasive adjunct to current AS strategies in PCa. mpMRI = multi-parametric magnetic resonance imaging, PSA = prostate-specific antigen.

Differential gene expression (DGE) analysis between RP and SD groups identified three significantly up-regulated genes *MT-TM* and *MT-TV*, which encode mitochondrial tRNAs (mtRNAs), and the unannotated *ENSG00000279123* enriched in RP patients, and one significantly down-regulated gene, *PLBD2* (Fig. 1C-E, Supp. Fig. 1C). Cell-of-origin analysis indicated that most cfRNA originated from non-tumoral sources, including smooth muscle, stromal, and immune cells (Fig. 1F), while epithelial signals were low (Supp Fig. 1D).

The small number of differentially expressed genes likely reflects the limited RNA input and modest cohort size typical of feasibility studies, and therefore these findings should be interpreted with caution. The low epithelial-cell enrichment scores may also contribute to the limited number of differentially expressed genes observed in patients with recurrent disease under AS, as epithelial cells are a known source of tumour-derived cfRNAs in PCa^[4]^. Purifying extracellular vesicles prior to cfRNA extraction could enhance detection and improve the sensitivity of RNA detection in liquid biopsy approaches^[4]^. The mtRNAs identified may represent useful biomarkers despite no difference in the overall mitochondrial cfRNA fraction between groups (Supplementary Fig. 1E) given that mitochondrial RNAs are stable in circulation^[3]^ and dysregulated in prostate cancer^[5]^. Read length and alignment rates differed significantly between RP and SD patients, indicating broader differences in cfRNA sequencing profiles.

In conclusion, our findings support the feasibility of transcriptomic profiling in PCa patients on AS. Future studies incorporating larger, longitudinal cohorts and integrating cfRNA signatures with established genomic classifiers could help determine whether cfRNA profiling can be used as a minimally invasive adjunct to current AS strategies in PCa (Fig. 1G). The ultimate goal is to improve baseline risk stratification and aid monitoring to guide personalised AS schedules.

## Acknowledgements

The authors would like to acknowledge the work of Daniela Barreto, the Clinical Research Coordinator, for preparation of slides and storage and preservation of plasma samples, the Garvan/St Vincent’s Prostate Biobank for storage and management of prostate tissue and blood samples, and the Garvan IT Applications Group.

## Conflict of interest

PR has received reimbursement for consultancy and advisory boards from Antev Ltd and Medtronic Ltd, speaker fees from Janssen-Cilag Ltd (Johnson & Johnson), and educational support from Angiodynamics UK Ltd, EDAP TMS S.A., HC 21 Healthcare Ltd, Medtronic Ltd, and Janssen-Cilag Ltd (Johnson & Johnson). JGF has received reimbursement for consultancy from Baseimmune Ltd (UK).

## Funding

This work was funded by the John Black Charitable Foundation and the Urology Foundation (to PR), Barts Charity (MGU0533 to PR and JGF), Orchid Charity, and Cancer Research UK Barts Centre Development Fund (to PR). PR was supported by a joint Royal College of Surgeons of England/Cancer Research UK Clinician Scientist Fellowship in Surgery (C19198/A15339). The Cancer Institute of New South Wales and St Vincent’s Prostate Research Cancer Centre funded the maintenance of the prostate biobank and CANSTO database.

## Data availability

Original fastq files will be shared via NCBI Gene Expression Omnibus (GEO).

## Authors’ contributions

JGF and PR conceived of and designed the study. JGF, EW, CAM, JT, PS and A-MH acquired and analysed the data; JGF performed statistical analyses. All authors interpreted the data. JGF and PR wrote the manuscript and all authors reviewed it. JGF and PR obtained funding.

## Figure Legends

**Supp. Fig. 1 - Additional analysis of cfRNA sequencing**

(A-B) Violin plots representing: (A) average read length (B) percentage of reads mapping to the hg38 genome. Comparisons are between SD and RP patients, Mann-Whitney U test, ** p<0.01, *** p<0.001.

(C) Principal component analysis (PCA) of variance stabilised read counts, colour of dots indicates progression status (SD = Stable Disease, RP = Progressed to Radical Prostatectomy). (D-E) Violin plots representing: (D) xCell cell epithelial cell type enrichment score derived from gene expression matrix (TPM) and (E) the fraction of mitochondrially encoded transcripts (calculated from TPM). Comparisons are between SD and RP patients, Mann-Whitney U test, not-significant (ns) p>0.05.

