## Supplementary figures and images for "Transcriptome profiling of cell-free RNAs in prostate cancer patients on active surveillance"

### Supplementary Figure 1

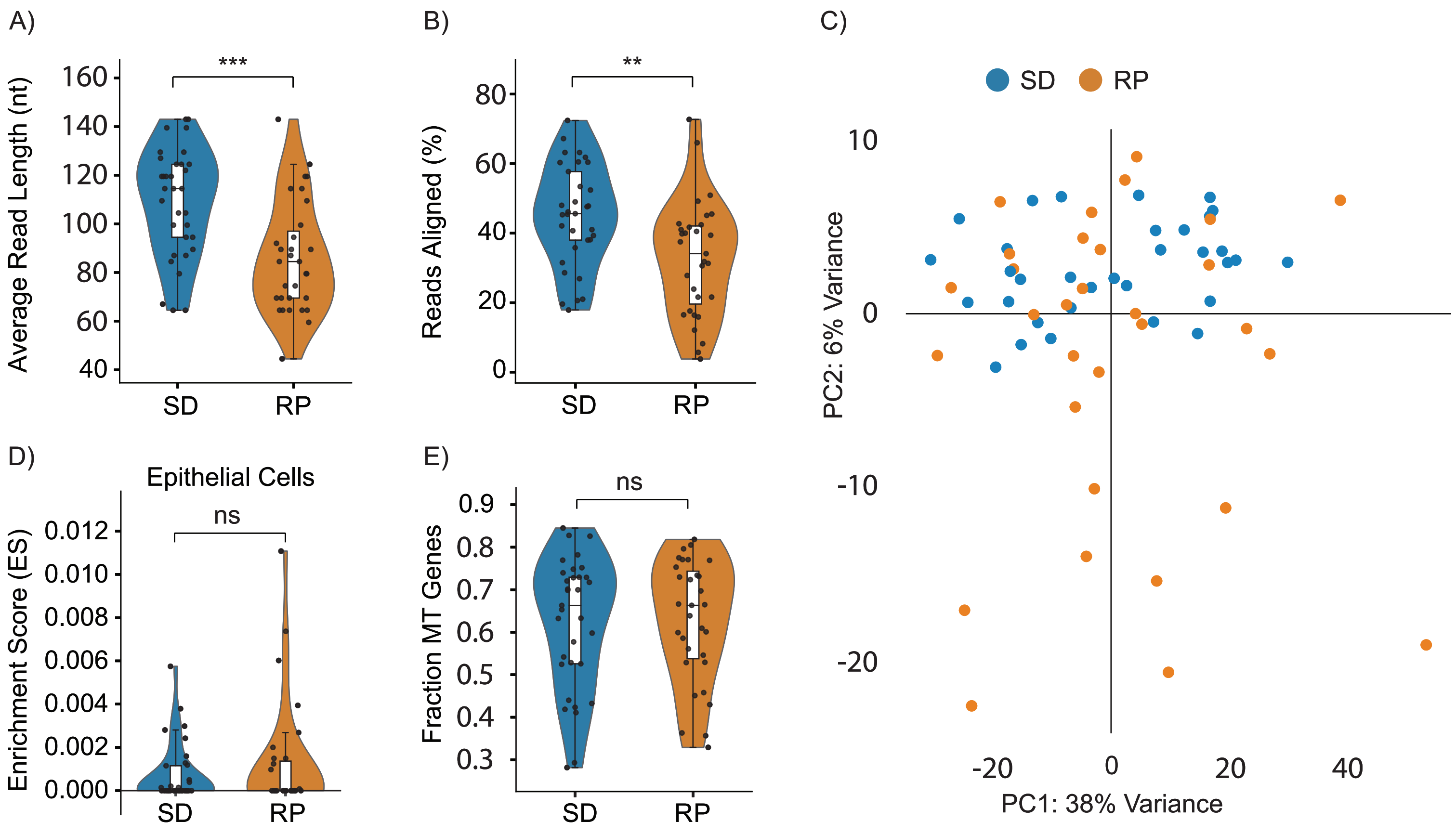
