## Supplementary Methods for "Transcriptome profiling of cell-free RNAs in prostate cancer patients on active surveillance"

### Sample collection and ethical approval

The Australian Prostate Cancer BioResource (APCB) collected biological samples and clinical data from men diagnosed with and undergoing treatments for prostate cancer at the Royal Adelaide Hospital (RAH) and St Vincent's Hospital (SVH), Sydney. All men gave written consent for sample and ongoing data collection, and practices and procedures were approved by the RAH and SVH Human Research Ethics Committees. Blood was collected according to the APCB's standard operating procedures, which have been reviewed within Australia and by the International Society of Biological and Environmental Repositories. Samples were collected in 8-ml Serum Separator Gel Vacuette tubes (Greiner BioOne, Frickenhausen, Germany; catalogue number 455078), allowed to clot for ~30 min at room temperature and centrifuged at room temperature for 10 min at 2500 g. Serum was aliquoted into Nunc cryovials (ThermoFisher Scientific, San Jose, CA, USA), snap frozen in liquid nitrogen and stored at −80 °C. Patients were followed up by clinicians typically at 3-month intervals with a PSA test. The study received ethical approval (Ref: 2019/ETH02834: Clinical, pathological and molecular markers of disease aetiology and outcome in prostate cancer and prostatic diseases).

### Sample processing and RNAseq library preparation

Circulating nucleotides were extracted from patient serum samples using QIAamp ccfDNA/RNA Kit (QIAgen, 55184) according to manufacturer's instructions. cfDNAs were removed by DNAse treatment, preserving cfRNA for downstream analysis. Libraries were prepared using SMARTer® Stranded Total RNA-Seq Kit v3 -Pico Input Mammalian (Takara, 634487) according to manufacturer's instructions. This includes an rRNA and mt-rRNA removal step and addition of an 8 nucleotide unique molecular identifier (UMI) for downstream read deduplication. Libraries were run on a NovoSeq 6000 with 150bp paired-end settings by Source BioScience.

### RNAseq read processing, alignment and counting

Paired fastq reads were trimmed and adaptors removed with fastp version 0.24.0 with a Quality Phred score cutoff of 15 and filtering of reads <30 bases. Umis (N) and Umi linkers (C) were extracted using umi_tools version 1.1.6 with pattern NNNNNNNNCCCCCC and appended to the read headers. Processed reads were mapped to human genome hg38 GCA_000001405.15 index using Hisat2 version 2.2.1 in paired end mode. Reads in Bam files were deduplicated using umi_tools version 1.1.6, discarding unpaired and chimeric reads, and sorted and indexed using samtools 1.21. Read mapping statistics were extracted from deduplicated Bam files using qualimap version 2.2.2a. Deduplicated reads were feature annotated and counted using featureCounts from subread version 2.0.8 and gencode v49 basic annotation gtf file.

### Gene expression analysis

Differential gene expression (DGE) analysis was performed using DESeq2 version 1.46.0 in R version 4.4.1 and RStudio. Log2 foldchange, pvalues, and padj values were calculated using the formula: lfcShrink(dds, coef="Group_rp_vs_sd", type="apeglm")[1]. Gene Ontology analysis was performed on a relaxed set of DGE using clusterProfiler version 4.14.6 with the function gseGO(geneList = gene_list, OrgDb = org.Hs.eg.db, keyType = "ENSEMBL", ont = "ALL", minGSSize = 10, maxGSSize = 500, pvalueCutoff = 0.05, verbose = FALSE).

**TPM Calculation**

Transcript abundance in Transcripts Per Million (TPM) was calculated by dividing raw gene counts by gene length in kilobases obtained from the featureCounts output, followed by scaling by the sum of length-normalised counts per sample and multiplying by $10^6$.

**xCell Analysis**

Cell type enrichment scores (ES) were calculated at the sample level from transcript abundance (TPM) using the xCellAnalysis function from xCell version 1.1.0 in R version 4.4.1 and RStudio[2]. Average ES for each cell type was calculated by taking the mean value for each condition (SD and RP). Heatmap filtered by cell type with mean ES > 0.05 was plotted using pheatmap version 1.0.13 in R version 4.4.1 in RStudio.
